# Kin-number distributions over age, sex, and time

**DOI:** 10.64898/2025.12.08.692903

**Authors:** Joe Butterick

## Abstract

Mathematical kinship demography is an expanding area of research. Most models explore the expected number of kin without accounting for demographic stochasticity. Recently, a paper provided a method to calculate the complete number-distribution of kin in a one-sex time-invariant demography. We extend this method to the case of two-sexes and to time-variant demographic rates.

Drawing from the mathematical tools of Fourier and convolution theory as well as basic probability and matrix algebra, we derive closed form expressions which capture the recursive nature of kin replenishment, generation-by-generation. Formulae presented here extend arbitrary genealogical distances to recover relatives considered in the leading frameworks of kinship. All we require as inputs are age, sex, and time-specific mortality and fertility schedules.

This research presents the first kinship model able to predict the probable numbers of relatives, structured by age and sex within a time-varying demography. As well as producing the probable numbers of living kin, the model flexibly extends to give the probable numbers of deaths an individual experiences. Such a detailed analysis of the kin-network will be useful in many fields.

## 1 Introduction

Understanding how the presence or absence of relatives affects an individual’s social environment is a fundamental question, focusing research in fields ranging from ecology to human demography. Kinship structure plays a considerable role in shaping the life-history evolution of many species. Interactions between kin may be within or between generation, and can facilitate benefits of increased protection (Clutton-Brock et al., 2001), the transfer of information (Ellis et al., 2024), resources (Watcher, 1997), and social learning (van Schaik, 2010; Thornton & Clutton-Brock, 2011), but to name some. Moreover, the theory of kin-selection^1^ (Hamilton, 1964) in animal populations remains one of the principal explanations for the evolution of cooperation (e.g., see Nowak (2006)). In humans, family plays a crucial role in supporting individuals. Recent demographic trends have been characterised by increasing lifespans and lower fertility, resulting in smaller family units in which multiple generations overlap at any one time. As such, a so-called “sandwich generation” of middle-aged individuals has consequently emerged. The sandwich generation must concomitantly support grand-children (thus enabling children to work), and parents – who on average being elderly also require care (Alburez-Gutierrez et al., 2021). Irrespective of whether ecological or human context, the number of kin an individual has available and can thereby interact with, depends on the reproduction of the individual, their ancestors, and their descendants.

The ever expanding field of mathematical kinship demography attempts to quantify such availability of kin. Models of kinship, emanating from Goodman et al. (1974), have recently diverged in two broad directions, one emphasising a greater focus on human kin (Caswell, 2019, 2020; Caswell & Song, 2021; Caswell, 2022, 2024; Butterick et al., 2025a) and another of more general ecological scope (Coste et al., 2021). With the exception of Butterick et al. (2025b) and Caswell (2024), in most of these models, kin availability has been measured through the “expected number”. Though projecting the mean numbers of kin contributes considerable advances in theoretical knowledge, it does not capture how demographic stochasticity – the inherent randomness of birth and death processes – affects kinship dynamics. Moreover, knowing the distribution for the number of kin offers valuable information, e.g., the further moments of, and variance in, kin-number.

Caswell (2024) proposed the first model to investigate stochasticity in kin, allowing one to obtain variances in kin-number. The ingenious method projects kin through a multi-type Galton-Watson branching process in matrix form, whereby age-classes are treated as types. Pioneering research by Coste (2025b) also holds great promise in theoretically explicating the role of stochasticity in kin. In a mathematically concise and elegant manner, the author utilises so-called genealogical Markov chains to project kin structured by generation number (or “trait”). Stochastic extensions of the work, again, could appeal to multi-type branching processes. Such progress will provide valuable tools for obtaining the number distribution of kin, especially in populations of arbitrary structure. Perhaps a more natural way to explore the probabilistic nature of kin accumulation, is through micro-simulation. This method allows one to obtain the exact number distribution (Wachter et al., 1997, 1978; Hammel, 2005; Margolis & Verdery, 2019), yet at the expense of computational challenge. This restriction on the applicability of simulating kinship networks (notwithstanding recent work in by Thiele et al. (2023)) makes analytical models highly desirable.

Tuljapurkar et al. (2020) proposed the first model – based on convolutions and Fourier transforms – to derive the distribution for an individual’s so-called lifetime reproductive success (i.e., the number of offspring produced over its lifetime). Drawing from similar mathematical tools, the first analytical model to calculate the exact number-distributions of generic kin was presented by Butterick et al. (2025b). The authors provide the probabilities that a typical population member – referred to as Focal (a convention adopted in the leading models of Caswell and henceforth) – will experience a given number of a certain kin-type, by age of Focal and by age(s) of the kin. That model operates in an age-structured one-sex time-invariant demography. In this research, we extend the framework to operate within a two-sex population subject to time-variant vital rates. We additionally propose a novel method for calculating the probabilities that an individual will experience a given number of deaths of a certain kin-type throughout their life-course.

In what follows, we seek to find, for each age of Focal *y* and corresponding time period *t*, the probability that she (or he) will experience an integer number *j* of a certain kin-type of given age and sex. In particular, for arbitrary kin-type “*k*”, we propose a block-structured vector of probability mass functions

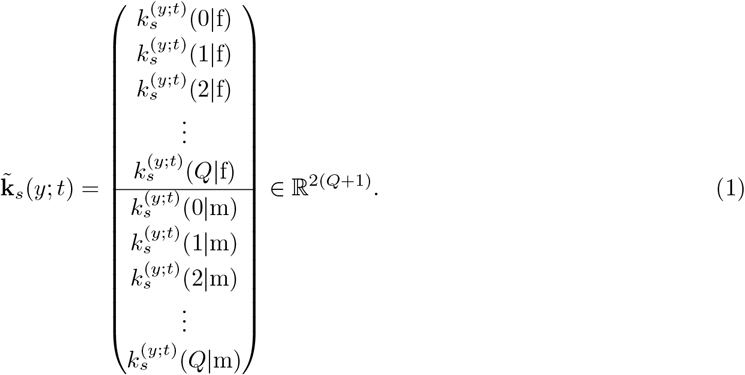

In Eq (1), the (*j* − 1)-th entry, 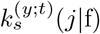, gives the probability that Focal of age *y* at time *t* has *j* female of that kin-type of age *s*. The (2(*j* − 1) + 1)-th entry yields analogous results for males of that kin-type. The variable *Q* defines the maximum lifetime kin-number for each kin-type of specific sex (e.g., aunts), and thereby, 2*Q* bounds the total number of each kin-type (e.g., aunts and uncles). Henceforth, from a notational perspective, we will consistently index kin-number through *j* and kin-sex through *γ* ∈ {f, m} (sometimes subscripted to distinguish from sex of producer kin and newborn kin). Moreover, we encode numerically the alphabetic indexes *γ* = f → 1 and *γ* = m → 2 for use in analytical derivations.

### 1.1 Notation

Stochastic matrices are denoted using blackboard bold – 𝔸, other matrices are denoted boldface upper-case **A**, vectors lower-case boldface **a**, and distributions are boldface Greek symbols, e.g., ***ψ***. When possible matrix entries will be given by lower-case letters, e.g., *a*_*i,j*_ the *i, j* entry of **A**, but when notation is a pain – for instance when the matrix is a function of parameters; **A**(*x*), we may use [**A**(*x*)]_*i,j*_. The transpose of a matrix is denoted by *†*, the Kronecker product of matrices by ⊗, the vec-operator vec(.), the Hadamard product ⊙, and the Hadamard product of a sequence of matrices is written:

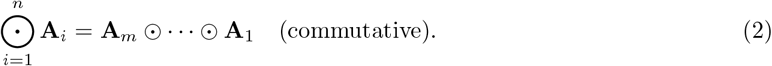

Being aware of the Hadamard notation above, we denote function composition by (*f* ∘ *g*)(*x*) = *f* (*g*(*x*)) and moreover, *f* ^[*n*]^(*x*) = (*f* ∘ … ∘ *f* )(*x*) as the n-times composition of *f*, and the ordered composition of functions {*f*_*i*_} by

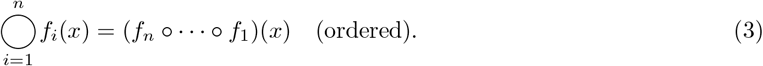

A discrete convolution of two functions *f* and *g* defined on the integers is given by 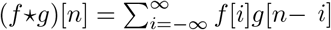. For two distributions ***ψ***_1_ ∈ ℤ^*n*^ and ***ψ***_2_ ∈ ℤ^*n*^ we write the discrete convolution as ***ψ***_1_ *\** ***ψ***_2_ with *m*-th entry defined through 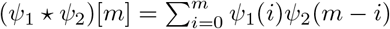. Over distributions ***ψ***_*j*_, …, ***ψ***_*n*_, we write

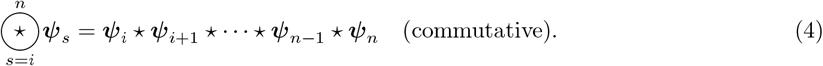

To represent the *n*−th convolution power of a distribution, we write

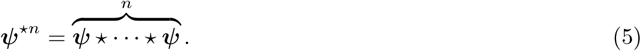

The discrete Fourier transform (DFT) for a distribution ***ψ*** = (*ψ*(0), …, *ψ*(*N* )) is defined through

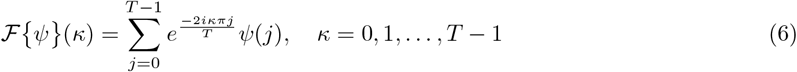

where 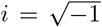, the *κ*-th entry is encoded by frequency *κ/T* for integer *T*, and *ψ*(*j*) = 0 for *j* > *N* + 1 (the output is zero padded at lengths greater than the input data). A useful mathematical equivalence, frequently applied henceforth, is that the inverse Fourier transform of the element-wise product of *m* Fourier transformed distributions ***ψ***_*j*_ where *j* = 1, …, *m* equals the convolution of the distributions:

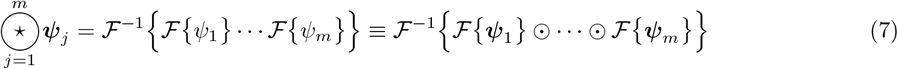

## 2 The model

The model closely resembles the one proposed in Butterick et al. (2025b). Focal and kin are once again defined through a common ancestor (Pullum, 1982), however, we now anchor the ancestor in time period of, as well as age of, their reproduction. Kin replenishment follows the same outline as the original model, but now with two-sex reproduction and time-specific fertility schedules. Survival extends both sexes and depends on time period.

### 2.1 Defining kin: Focal’s *q*-th ancestor and their *g*-the descendant

Suppose that Focal is of age *y* at time *t*. Let *q* be the number of generations separating Focal and the nearest common ancestor which relates them to their kin. Let *g* the number of generations separating Focal’s kin from this ancestor. Let *b*_*i*_ represent the age at which Focal’s *i*-th generation ancestor gives birth to Focal’s (*i* − 1)-th generation ancestor. Define 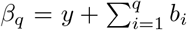 representing the age of Focal’s *q*-th ancestor when Focal is *y* (at time *t*). Here we only talk of female ancestral reproduction; we adopt the female dominance assumption of Caswell (2022). Define *s*_*i*_ as the current age of Focal’s kin (at time *t*) related to Focal as the *i*-th generation descendant from Focal’s *q*-th ancestor. Thus, the age at which Focal’s (*i* − 1)-th generation descendant produces Focal’s *i*-th generation descendant is *s*_*i*_ − *s*_*i*+1_. Lastly, define n as the minimum and 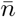 the maximum fertile age (over both sexes). Let the vector of probability mass functions for Focal’s *g, q* kin, who can be of age within some feasible range, *s*_*g*_ ∈ Σ, when Focal is age *y* at time *t*, be denoted by 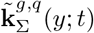. For any sub-interval *J* ⊆ Σ, we recover 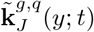, and age-specific pmfs are denoted 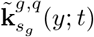.

### 2.2 Survival

Survival is independent between sexes, but depends on age-and-time specific mortality rates. Thus, we create sex-and-time-dependent analogues of 𝕌(*s*^′^, *s*) defined in Butterick et al. (2025b). For kin of sex *γ*, let the matrix 𝕌^*γ*^ (*s*^′^, *s*; *τ* ) act on a kin-number pmf representing kin of age *s* at time *τ*, and project these number-probabilities of the kin, conditional on survival, through to the kin being age *s* at time *τ* +(*s* − *s*^′^ ). To ensure these matrices act independently on the sex-structured pmfs, we apply 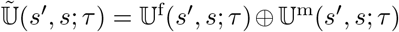.

To construct each matrix, let *u*_*x,τ*_ (*γ*) be the probability that an individual of sex *γ* survives from age *x* at time *τ* to age *x* + 1 (at time *τ* + 1). Introducing the probability 𝒰 ^(*γ*)^(*j, l, s*^′^, *s, τ* ) that out of *j* kin of sex *γ*, aged *s*^′^ at time *τ*, some *l* survive to age *s*:

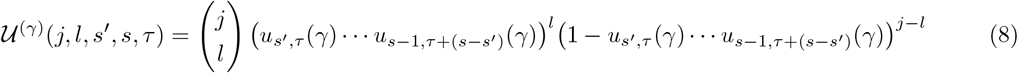

we define matrix entries through

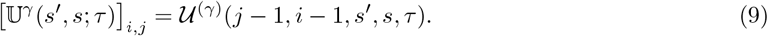

See Butterick et al. (2025b) for further details.

### 2.3 Reproduction

Define the probabilities that an individual of sex *γ* and age *s* at time-period *τ*, has *j* = 0, 1, 2, …, *Q* female or male offspring, as a function of time-dependent vital rates:

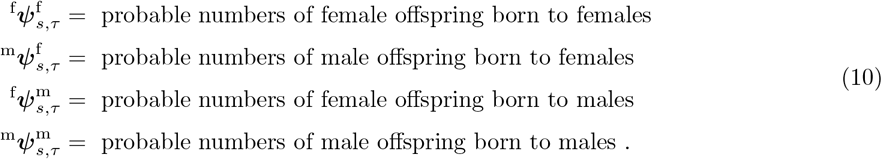

Using the convolution power method of Butterick et al. (2025b), define the matrix which projects a pmf of producer kin of sex *γ*_*p*_ ∈ {f = 1, m = 2} (“p for producer”), onto a pmf of newborn kin of sex defined by *γ*_*o*_ ∈ {1, 2} (“o for offspring”):

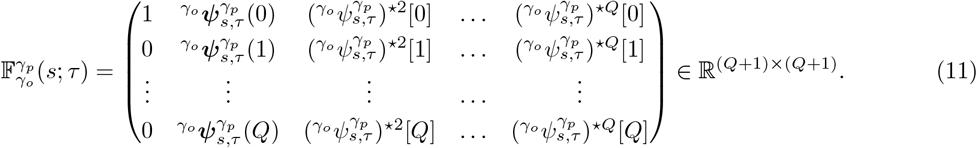

There are four distinct matrices 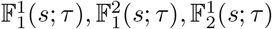, and 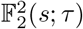 representing respectively, the births of: female newborns to female kin, female newborns to male kin, male newborns to female kin, and male newborns to male kin.

### 2.4 The probable ages of ancestral reproduction

We seek 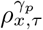: the probabilities that a newborn at time *τ* was born to a *γ*_*p*_ -sex parent of age *x* (i.e., 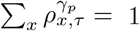). Introduce time-and-sex-specific survival 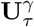 and fertility 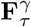 matrices which comprise, for each time period *τ*, the block-structured projection matrix

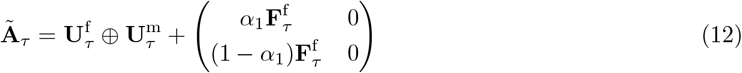

where *α*_1_ is the probability that a newborn is female. Let *τ* = 0 define the earliest set of demographic rates. Assume before this time, a steady-state demography with stable population structure 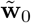. Set 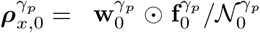 where 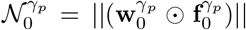, and 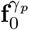 is the first row of 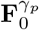, and 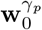 is the sex-specific extraction of the stable population structure. At *τ* = 0, the probability that the mother of a newborn is aged *x* is 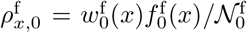; the probability that the father is aged *x* is 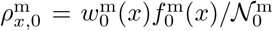 For each subsequent year in the time-variant situation, project

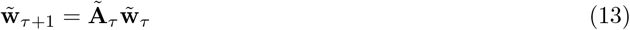

and apply the same procedure to obtain 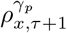 for all subsequent times *τ* ≥ 0 whereby vital rates vary.

### 2.5 The recursive structure of kin replenishment

Here we apply to the naturally recursive structure of kin replenishment; allowing for simplification of the mathematical exposition henceforth, through use of function composition. In principle, because each coming generation of kin is comprised of the reproduction of the previous generation, the *g*-th generation can be written as the composition of the reproduction of the *i* = 1, …, *g* − 1 generations. Section 2.5.1 and Section 2.5.2 respectively show how we implement generational reproduction in the cases of direct ancestors and collateral lineages.

#### 2.5.1 Ancestors

With our assumption of female reproductive dominance, first-generation descendants of a direct ancestor of Focal are produced by females only. As such, set

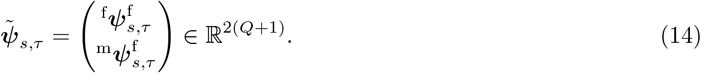

Since *β*_*q*_ = *y* + ∑ _*i*_ *b*_*i*_ is the age of Focal’s *i*-th ancestor at present, *t* − *β*_*q*−1_ represents the time-period at which Focal’s *q*-th ancestor produces Focal’s (*q* − 1)-th ancestor. The number of distinct *i*-th generation ancestors of Focal grows according to 2^*i*^. Encode each such ancestor by a sequence **a**_*i*_ = (*a*_1_, …, *a*_*i*_) where *a*_*j*_ ∈ {f, m}, *j* = 1, …, *i* indicates whether the ancestor of Focal’s (*j* − 1)-th generation ancestor is female or male. Basic examples are given in Appendix B and for a schematic see Fig 1. Define the set of unique ancestral sequences

**Figure 1.**
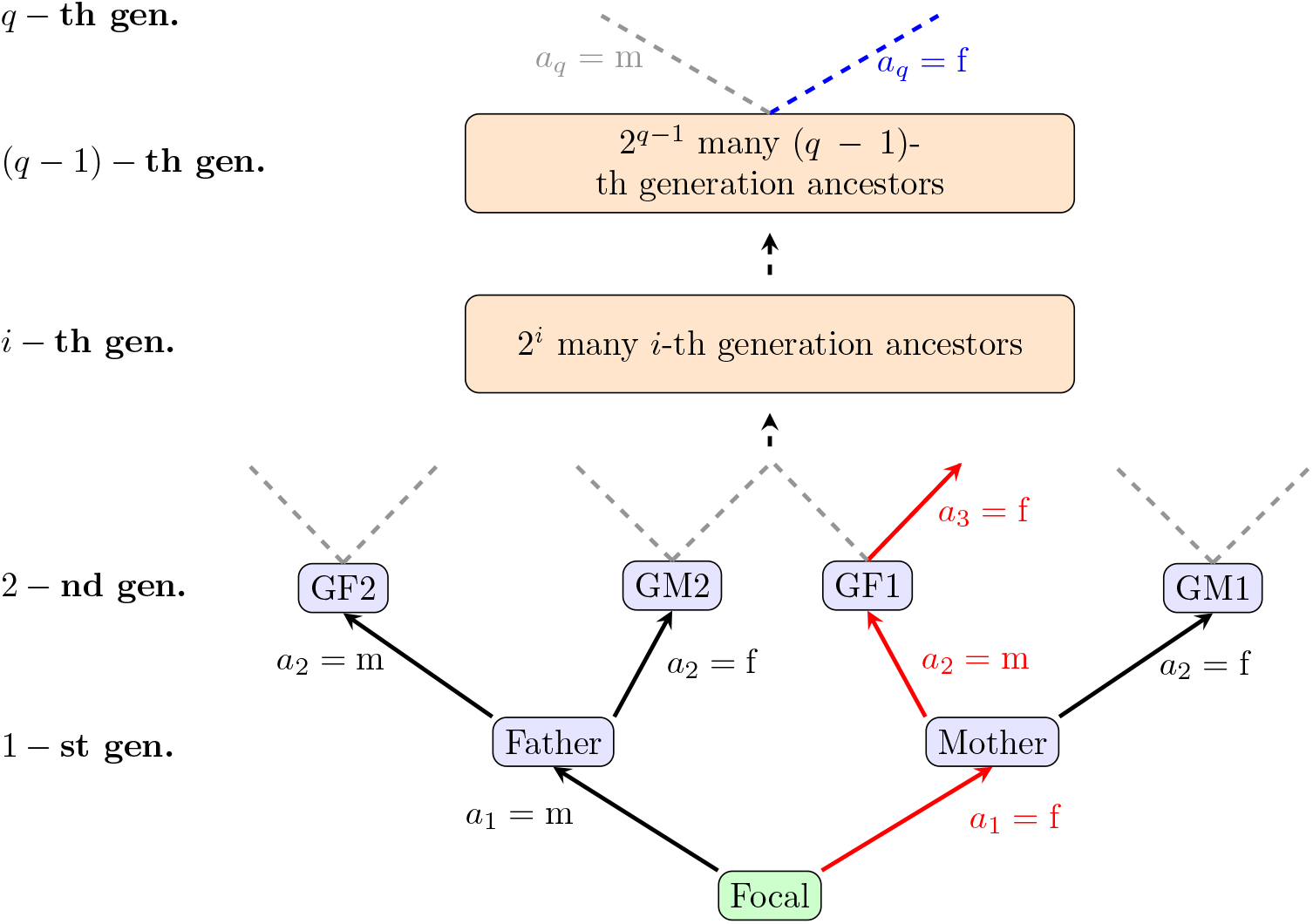
Ancestral sequences. Distinct ancestral sequences to Focal’s *q*-th ancestors are unique arrangement of *a*_1_, *a*_2_, …, *a*_*q*_. Each *a*_*i*_ = f,m represents the sex of the parent of Focal’s (*i* − 1)-generation ancestor. An example ancestral sequence, up to generation *i* = 3, is given in red. Each ancestral sequence is independent, and thereby, so are the lines of kin descendants emanating from each ancestor’s reproduction. The number of ancestral sequences to generation (*q* − 1), is equal to the number of routes from Focal to the generation, i.e., the size of the set 𝒞_*q*−1_. Starting from the *q*-th generation ancestors, we assume that only female ancestors reproduce (blue), resulting in 2^*q*−1^ independent lines of descendant kin.

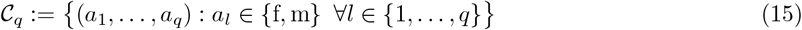

with cardinality |𝒞_*q*_| = 2^*q*^. We need only consider the 2^*q*−1^ female *q*-th generation ancestors, encoded through distinct sequences **a**_*q*−1_. Consider a particular *q*-th ancestor of Focal, defined by a distinct ancestral sequence **a**^*^ ∈ 𝒞_*q*−1_. Then for example, the vector of distributions for the number of female and male newborns (who would be age *s*_1_ > *β*_*q*−1_ when Focal is age *y*) of this particular ancestor aged *β*_*q*_ − *s*_1_ is:

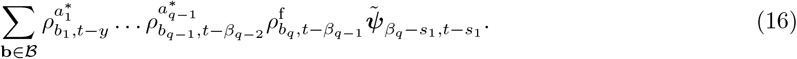

Above, for notational ease, we introduce the set

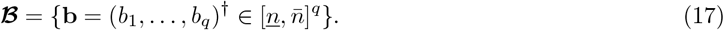

In Eq (16), while all routes back to Focal’s (*q* − 1)-th ancestor can go through male or female lines, Focal’s *q*-th ancestor is strictly chosen to be female.

#### 2.5.2 Non-ancestors

Consider at time *τ*, an arbitrary kin-type 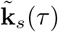 of age *s*. The random variables representing offspring numbers are independent between sex of producer kin. This means, for example, that the random variable for the sum of female offspring born to females and female offspring born to males is the convolution of the measures for the respective random variables. Hence, using Eq (11), the pmf for female offspring is 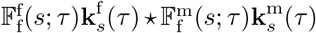. Using this information, recalling that we encode female (male) reproducer kin by *γ*_*p*_ = 1 (*γ*_*p*_ = 2) and female (male) offspring kin by *γ*_*o*_ = 1 (*γ*_*o*_ = 2), after a little algebra, we define the following operator which acts on an arbitrary (*i* − 1)-th generation descendant of Focal’s *q*-th ancestor:

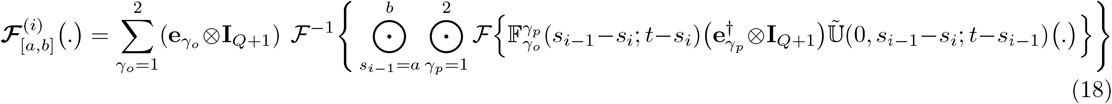

A detailed explanation of the above is given in Appendix A. Briefly put, Eq (18), through the action of 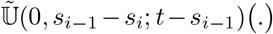. calculates the number of survivors of the arbitrary (*i* − 1)-th generation descendant of Focal’s *q*-th ancestor, from age 0 to age *s*_*i*−1_ − *s*_*i*_, and starting from time *t* − *s*_*i*−1_. Reproduction by this kin of female offspring is defined when *γ*_*o*_ = 1. Here, at age *s*_*i*−1_ − *s*_*i*_, one finds through the action of 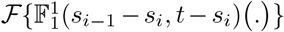. the DFT of the pmf of female newborns born to female kin (*γ*_p_ = 1), element-wise (Hadamard) multiplied through 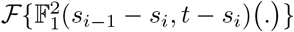, the DFT of a pmf of female newborns born to make kin (*γ*_*p*_ = 2). Taking the inverse DFT of this Hadamard product gives the overall pmf for female newborns, born to male and female kin of exact age *s*_*i*−1_ − *s*_*i*_. Taking the Hadamard product of 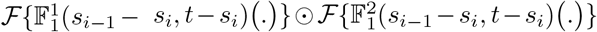, over fertile ages *a* − *s*_*i*_, …, *b* − *s*_*i*_, and then taking the inverse DFT, gives the pmf of female newborns born within the interval [*a* − *s*_*i*_, *b* − *s*_*i*_]. Call this **off**^fem^. Reproduction of the kin of male offspring is defined by *γ*_*o*_ = 2. Through the same procedure just described above, we obtain the pmf for male newborns (**off**^male^). The summation through (**e**_1_ ⊗ **I**_*Q*+1_)**off**^fem^ + (**e**_2_ ⊗ **I**_*Q*+1_)**off**^male^ stacks the two distributions resulting in the original block-structured vector of probability mass functions.

#### 2.5.3 Replenishment of kin

Given the information in Section 2.5.1 and Section 2.5.2 and assuming *q* > 0, while *g* = 1, the first-generation descendants of Focal’s *q*-th ancestor are strictly defined by female reproduction distributions. All subsequent generations of descent can be found by convolving the previous generation descendants’ possible reproduction distributions, over age and sex. This recursive nature of kin-replenishment allows us to write Focal’s [*g, q*]-kin as a composition of the operator in Eq (18), as we show below (see Section 2.6-Section 2.9).

### 2.6 Kin which descend through younger siblings of Focal’s (*q* − 1)-th ancestor

Using the operator in Eq (18) we define the cases:

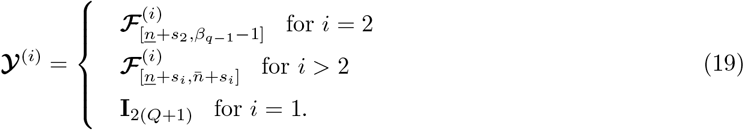

For each of Focal’s possible 2^*q*−1^ distinct female ancestors, uniquely determined by **a** ∈ 𝒞_*q*−1_, we set

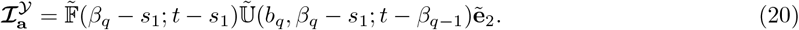

In Eq (20), the operation 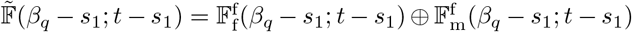 yields the reproduction of a particular *q*-th ancestor when the ancestor is of age *β*_*q*_ − *s*_1_ (at time *t* − *s*_1_). At this reproductive event the ancestor procured a younger sibling of Focal’s (*q* − 1)-th ancestor. Here, 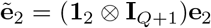 is a block structured vector of unit vectors, each with mass in the second entry – ensuring the ancestor is alive at the reproductive event.

Conditioning on ancestral sequence **a** ∈ 𝒞_*q*−1_ leads to a distinct ancestor. The reproduction of the ancestor is then conditioned on the respective probabilities that a specific set of parental ages, **b** ∈ ℬ, defines the demographic events resulting in their past reproduction. For each fixed **b**, the resulting kin-number random variable for Focal’s kin of exact age *s*_*g*_ is given by the composition 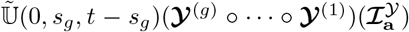. Due to independence in the parental age distribution, for fixed **b**, the number of Focal’s kin now of age in the range *s*_*g*_ ∈ Σ is the sum of the random variables representing kin-number of each age. The sum of these random variables is equal to the convolution of their probability measures (each given by a composition as above). The total number of kin which descend from each distinct *q*-th ancestor is therefore found by summing over all possible maternal ages **b**.

Due to independence in the ancestral sequences (see Fig 1 for illustration of independence), the total number of Focal’s kin is the sum of the random variables representing kin-numbers from distinct ancestors. Hence, total kin-numbers are found by the convolution of the probability measures defined by **a** ∈ ℬ_*q*−1_:

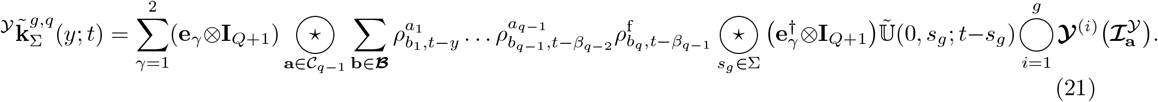

To summarise, Eq (21) provides the total pmf for the *g*-th generation descendants of Focal’s *q*-th ancestors, each of which descend through a younger sibling of one of Focal’s (*q* − 1)-th ancestors, and who now are of age *s*_*g*_ ∈ Σ. The feasible age-range Σ can be found in Appendix C.5. Age-specific pmfs are recovered by restriction to any scalar 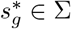.

### 2.7 Kin which descend through older siblings of Focal’s (*q* − 1)-th ancestor

Using the operator in Eq (18), define

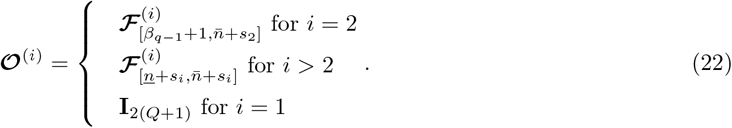

For each **a** ∈ 𝒞_*q*−1_ encoding one of Focal’s 2^*q*−1^ female *q*-th ancestors, define the reproduction of this ancestor at age *β*_*q*_ − *s*_1_, i.e., when they procured an older sibling of Focal’s (*q* − 1)-th ancestor:

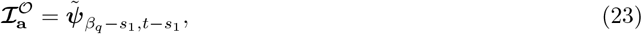

Using precisely the same arguments of independence and the application of convolutions when deriving Eq (21) of Section 2.6, we find the kinship formula for kin which descend through older siblings of (one of) Focal’s (*q* − 1)-th ancestors:

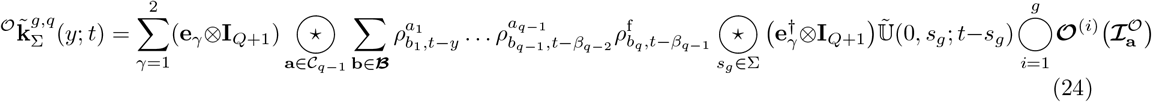

Eq (24) provides the total pmf for the *g*-th generation descendants of Focal’s *q*-th ancestors who now are of age *s*_*g*_ ∈ Σ (range found in Appendix C.5). Each of these kin descend through a older sibling of one of Focal’s (*q* − 1)-th ancestors. Examples are shown in Box 1. Age-specific pmfs are recovered by restriction to any scalar 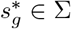.

Box 1

Suppose *n* = 1, 2, 3, 4 and reproduction in each age. Consider the number of nieces and nephews of age 2 or 3 (Σ = [2, 3]). We go up one generation to Focal’s mother, so 𝒞_0_ = {f}. Eq (24) becomes:

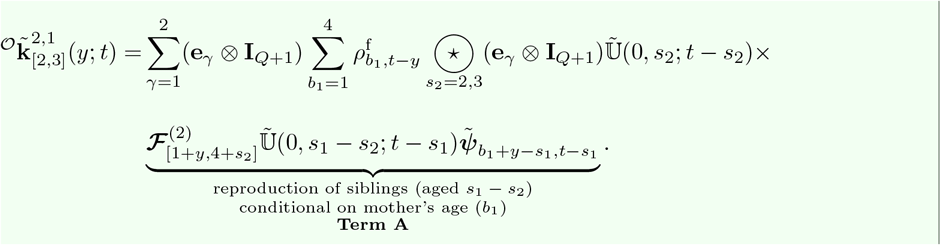

For example, condition on Focal’s mother being *b*_1_ = 1 at birth of Focal. **Term A** reads:

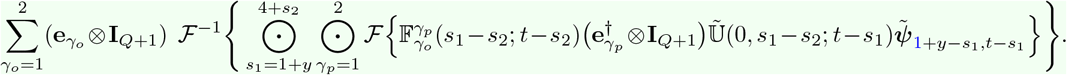

Consider nieces *γ*_*o*_ = 1. Suppose when Focal is *y*, nieces are *s*_2_ = 2. For *γ*_*p*_ = 1 and 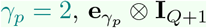 resp. selects Focal’s sisters’ and brothers’ reproduction:

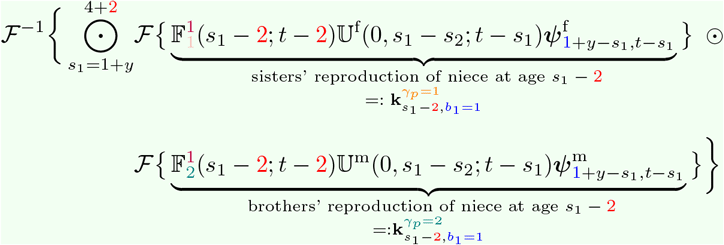

Expanding the Hadamand product over possible ages when siblings had nieces we have:

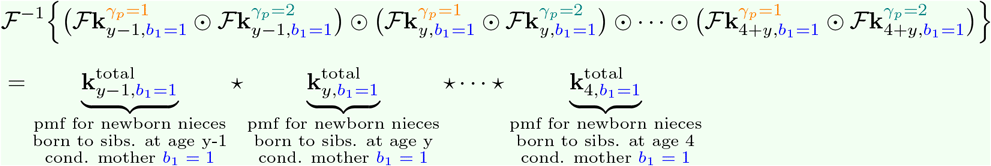

the pmf for newborn nieces, born to Focal’s siblings of all ages, and conditional on mother being *b*_1_ = 1 when having Focal. Nieces survive to age *s*_2_ = 2 with probabilities given in 𝕌^f^(0, *s*_2_; *t* − *s*_2_). The total number of nieces aged 2 or 3 is the sum of the independent age-specific random variables. This is equal to the convolution of the pmfs over 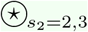.

The unconditional pmf is the weighted sum over the probable ages of mother. The same procedure applies for nephews.

### 2.8 Ancestors of Focal

Focal has up to 2^*q*−1^ many *q*-th ancestors of specific sex *γ* (see Fig 1). The plan to calculate the overall number pmfs, structured by sex, is as follows. Consider each sex separately. For each unique and distinct ancestor of given sex, we construct a pmf for its number distribution (with kin-number either zero or one). The (Binomial) random variables representing kin-numbers for each distinct ancestor are independent. We sum the random variables, which is equal to the convolution of the pmfs.

To simplify exposition we introduce the set

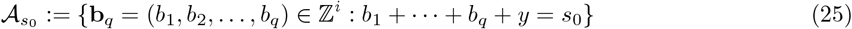

which counts the number of arrangements of genealogical ages of reproduction, with constraint that Focal’s *q*-th ancestor is aged *s*_0_ when Focal is *y*. For each distinct *q*-th ancestor of sex *γ*, the probability it is alive at present is

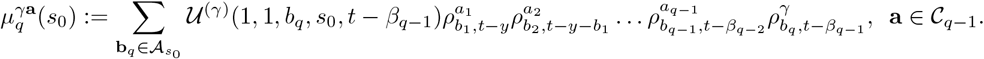

Here, *a*_*j*_, 1 ≤ *j* < *q* uniquely determines the line of ascent to Focal’s (*q* − 1)-th ancestor, while *γ* defines the sex of Focal’s (*q* − 1)-th ancestor’s parent; Focal’s *q*-th ancestor. The term 𝒰 calculates the probability that the ancestor survives from producing Focal’s (*q* − 1)-th ancestor up to age *s*_0_. Over the age-range Σ, which the ancestor can be when Focal is *y*, we find

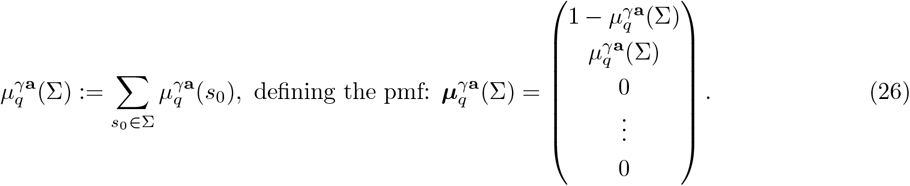

Eq (26) provides a probability number-distribution for a particular *q*-th generation ancestor of Focal over all possible ages, when Focal is *y*. Using the conditional independence of the kin-number random variables between distinct **a** ∈ 𝒞_*q*−1_, the total pmfs for Focal’s *q*-th generation ancestors (structured by sex) are:

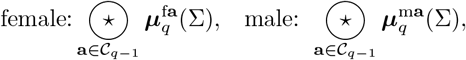

or, in block structured form:

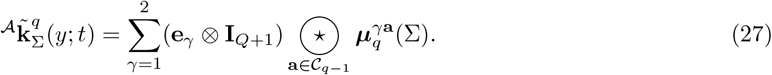

Age-specific pmfs can be readily obtained from the total pmfs simply by restricting to a scalar *s*^*^_0_ ∈ Σ to a scalar in Eq (26).

### 2.9 Descendants of Focal

Define the distribution to reflect the reproduction of Focal of sex *γ* and age *x* at time *τ*, by

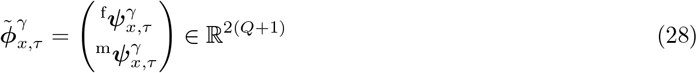

Focal’s first descendants are born to Focal, and have offspring pmfs defined by Focal’s sex, *γ*. Each subsequent generation of descent is born under two-sex reproduction. As such, define

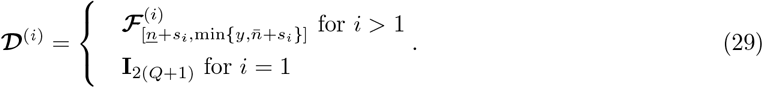

For Focal’s *g*-th generation descendants who can be of age *s*_*g*_ ∈ Σ when Focal is *y* at time *t*, we have the constraint 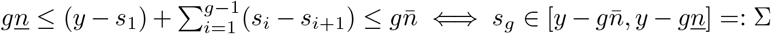. Then, the pmf for Focal’s *g*-th generation descendants is:

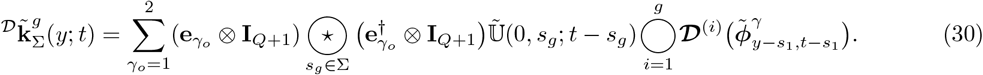

The random variables representing the offspring born to Focal and her offspring are independent. Thus, sex-specific convolutions over ages of descendants’ reproduction *s*_*i*_ − *s*_*i*+1_ yield the pmfs for newborns of the (*i*+1)-th generation. When *γ*_*o*_ = 1, Eq (30) extracts the pmf of Focal’s female *g*-th generation descendants of age *s*_*g*_, and stacks it on top of a zero vector of length *Q*. When *γ*_*o*_ = 2 the pmf for Focal’s male descendants is extracted and stacked below a zero vector of length *Q*. The sum of these two vectors is the resulting block-structured vector of pmfs. Restricting the feasible age range to a scalar 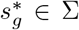 recovers the age-specific pmf.

### 2.10 Deriving the probabilities that Focal experiences *j* number of deaths of a kin-type

As well as providing the probable numbers of kin by age of Focal and age(s) of kin, our model also permits the probabilities of kin-loss. In Appendix C we show: (i) how to derive the unconditional probabilities that Focal experiences *j* many deaths of a kin by age *y* (here the probability of zero deaths includes the probabilities of kin never born). (ii) the conditional probabilities that Focal loses *j* many kin, given that the kin where born.

## 3 Application

Here, we use single year of age fertility and mortality estimates from 1938-2070 for England & Wales, historically sourced from the Office for National Statistics (ONS, 2024), and projected using median point estimates from the models of Hilton et al. (2018) and Ellison et al. (2023). In the data, male fertility rates go back to 1961. Thus we back-cast to 1938. A constant sex ratio of 105:100 males to females is assumed. We assume Poisson reproduction. We use Bernoulli mortality. *Q* = 10 is used as an upper-bound for sex-specific kin-numbers; that is, Focal is not assumed to experience more than 20 of any one kin-type. We assume that Focal is female here but could equally model the kinship of a male Focal.

### 3.1 Descendants

Fig 2 illustrates the probable numbers of sons and daughters that Focal has throughout her life, depending on which year she is born. Due to our forecast rates ending in 2070, the 2000 cohort of Focals live to age 70. We see that this cohort is predicted to have the lowest probability of having offspring. The 1940 cohort has the highest probability of having children: by age 50 of a typical Focal, she has a 0.37 chance of having one daughter and 0.20 change of 2 daughters; a 0.36 chance of having a son and 0.21 chance of two sons.

**Figure 2.**
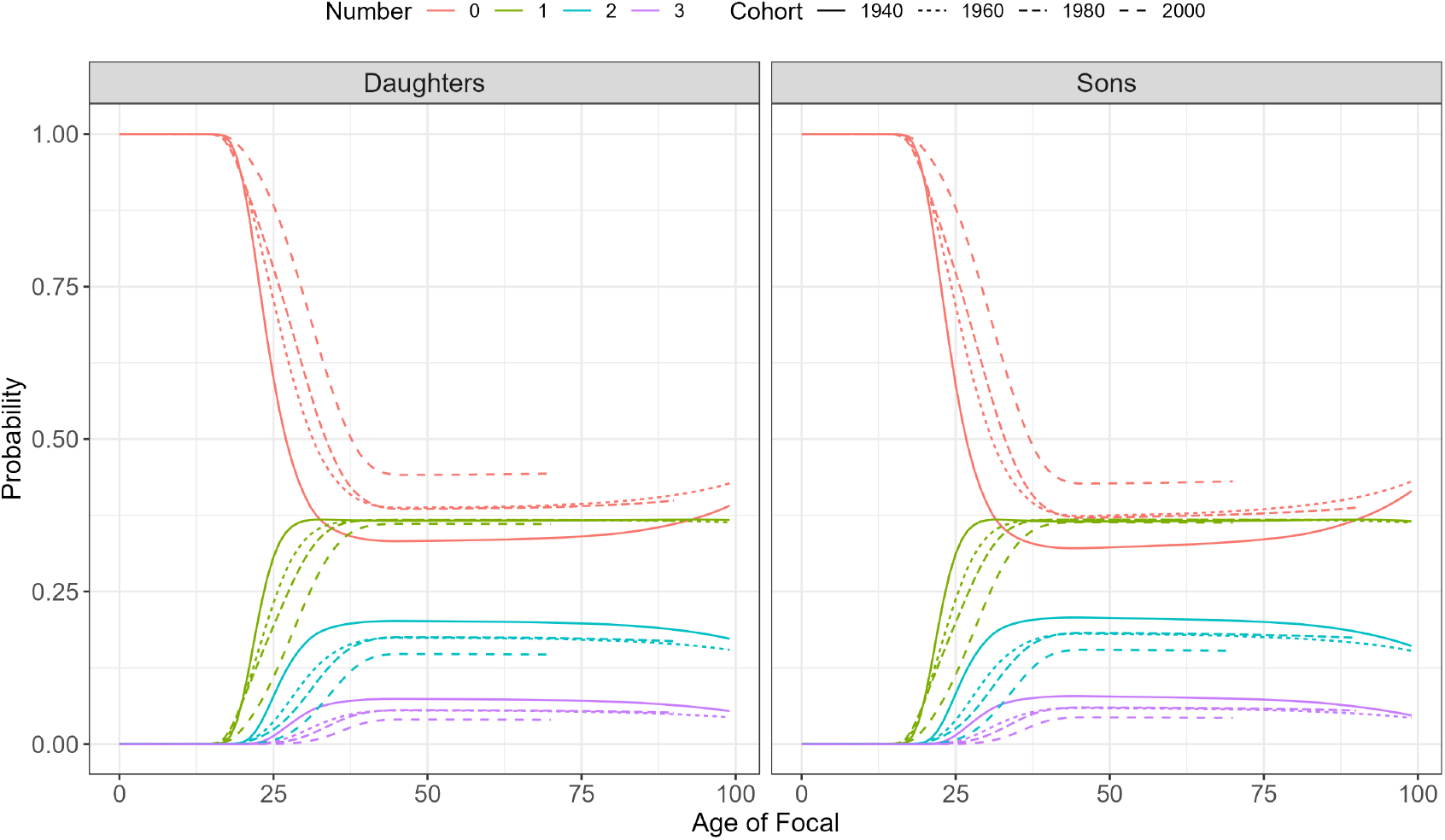
The probable number of offspring of Focal as a function of Focal’s birth cohort. Note that *j* = 4+ kin-numbers are not show for ease of interpretation.

### 3.2 Ancestors

#### 3.2.1 Parents

Fig 3 compares the probabilities that Focal has mother and father alive. The top row depicts these probabilities as a function of the cohort of birth of Focal, the bottom row as a function of the time period we sample Focal from. Improvements in mortality over time explain the monotonic increases in the age-specific probability of having an alive parent. Note that we here only project cohorts from 1940 to 2020: the 2050 cohort of Focal’s is not plotted and the 2050 period only includes Focals aged 30+.

**Figure 3.**
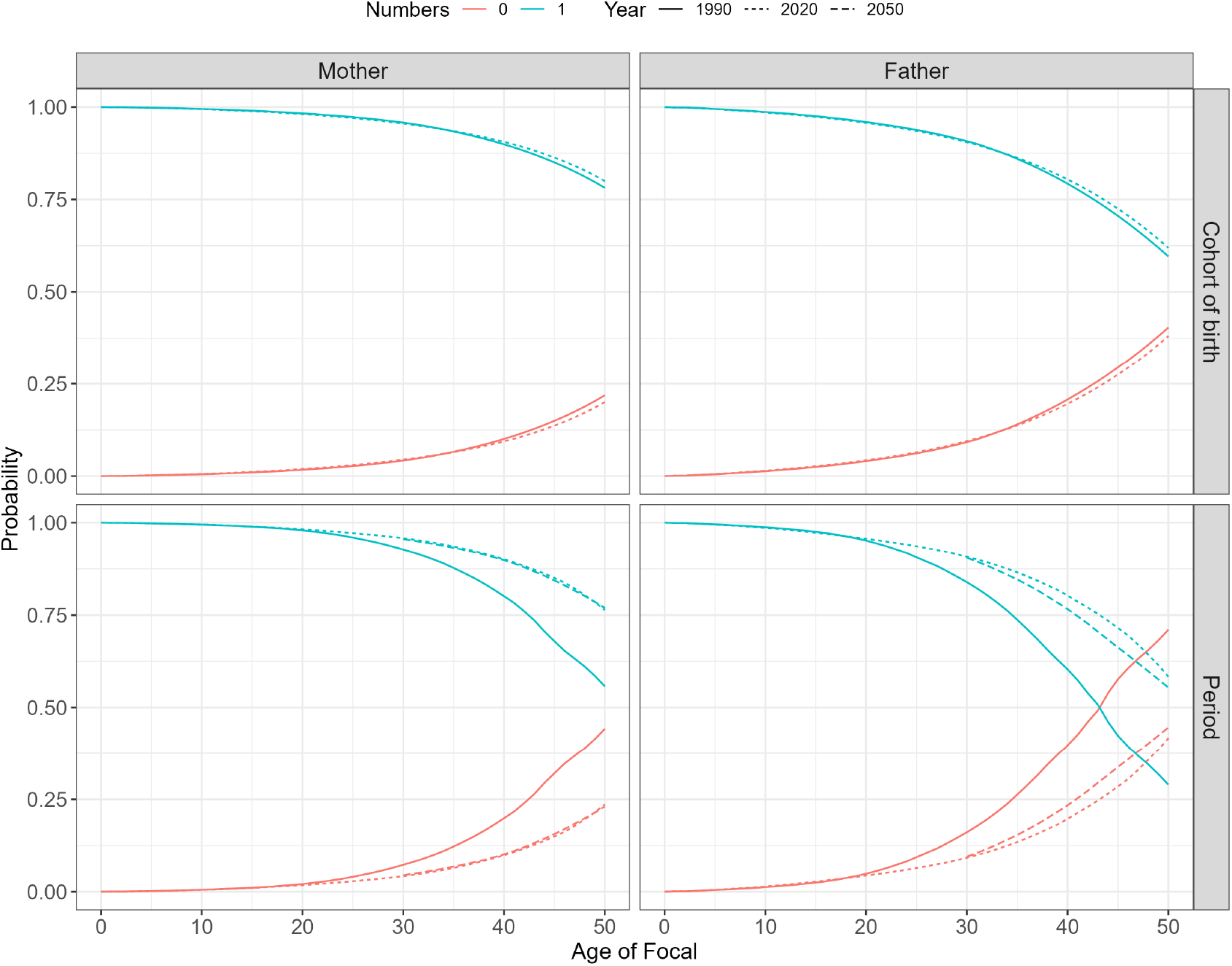
The probabilities (colour) the Focal’s parents are alive. Top: by age of Focals born to different cohorts. Bottom: by age of a typical Focal sampled from a given time period (here each age of Focal along the *x* represents a different cohort of Focals).

#### 3.2.2 Grand parents

Fig 4 compares, for Focals born to cohorts 1980, 2000, and 2020, the probable number of grandparents that she will have up to her being age 20. There are 4 possible grandparents (*q* = 2 and |𝒞_2_| = 2^2^), of which Focals born to the 2020 cohort are predicted to maintain more of. With probability 0.68, this cohort has all grandparents at birth. The 1980 cohort by contrast has all 4 grandparents at birth with probability 0.56.

**Figure 4.**
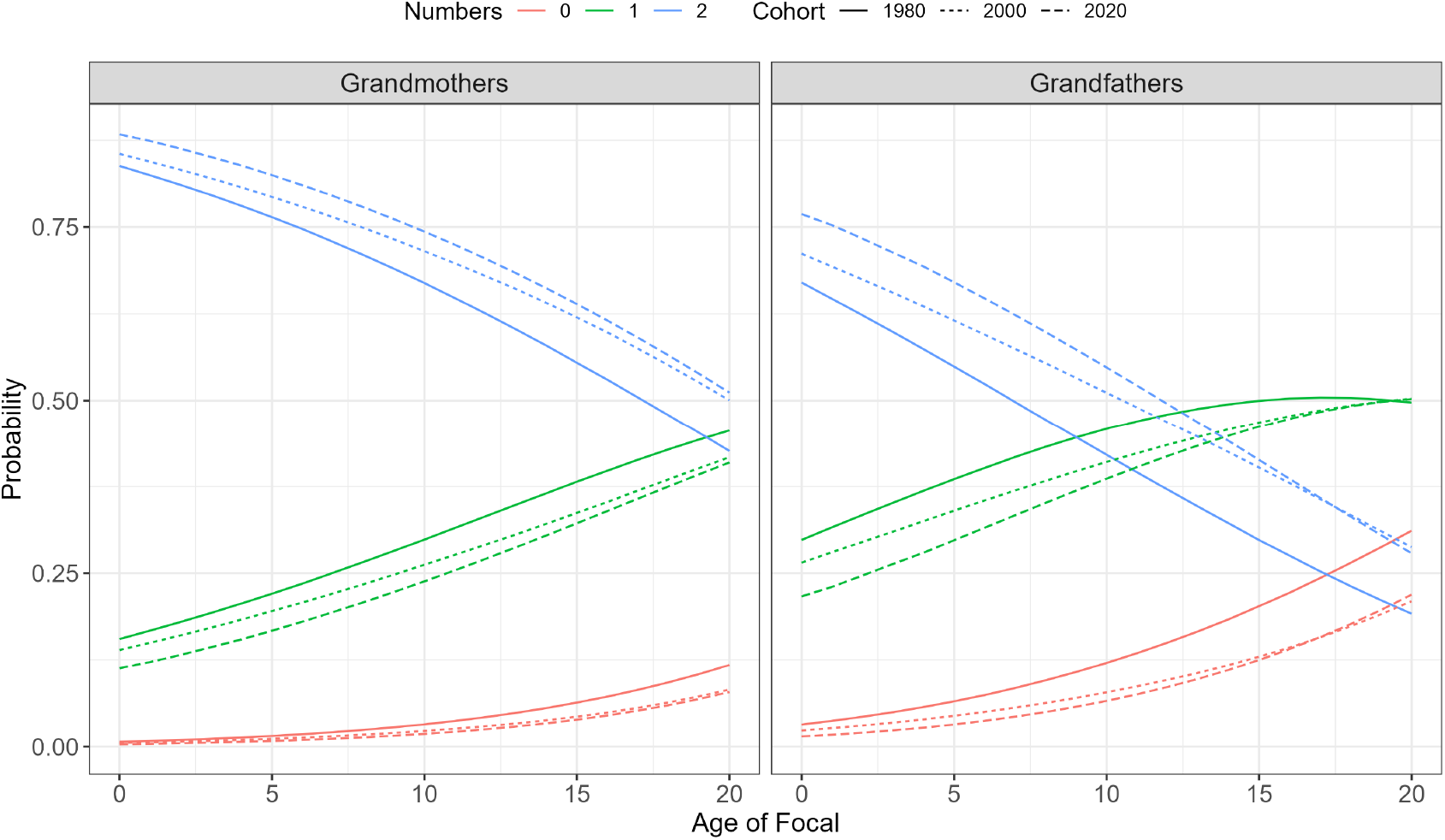
The probable number of Focals grandparents depending on her cohort. Colour shows number; line-type cohort; facet shows sex of relative. Note that here we run the model to Focal age 20 with time-variant rates from 1940-2070 (pre 1940 uses static rates).

### 3.3 Collateral kin

Fig 5 illustrates the total probabilities that Focal has *j* = 0, 1, 2, 3 siblings (over all ages the siblings can be), by age of Focal. We omit the *j* = 4+ probabilities for visual ease. The top row depicts these probabilities as a function of the cohort of birth of Focal, the bottom row as a function of the time period we sample Focal from. The 2040 cohort consist of 0-30 year old Focals due to our rate forecasts ending in 2070. The rich output allows for detailed analysis. One example is that the 1980 cohort of Focals are more probable to experience a sibling than any other cohort – a consequence of high fertility rates in the late 1970s and early 1980s compared to later times. Another example is the 1980 time period demonstrates an notable increase in the probability of not having a sibling from age 20 of Focal. This is likely due to the 20-40 year old Focals being born between 1940-1960 and having siblings which experienced higher mortality.

**Figure 5.**
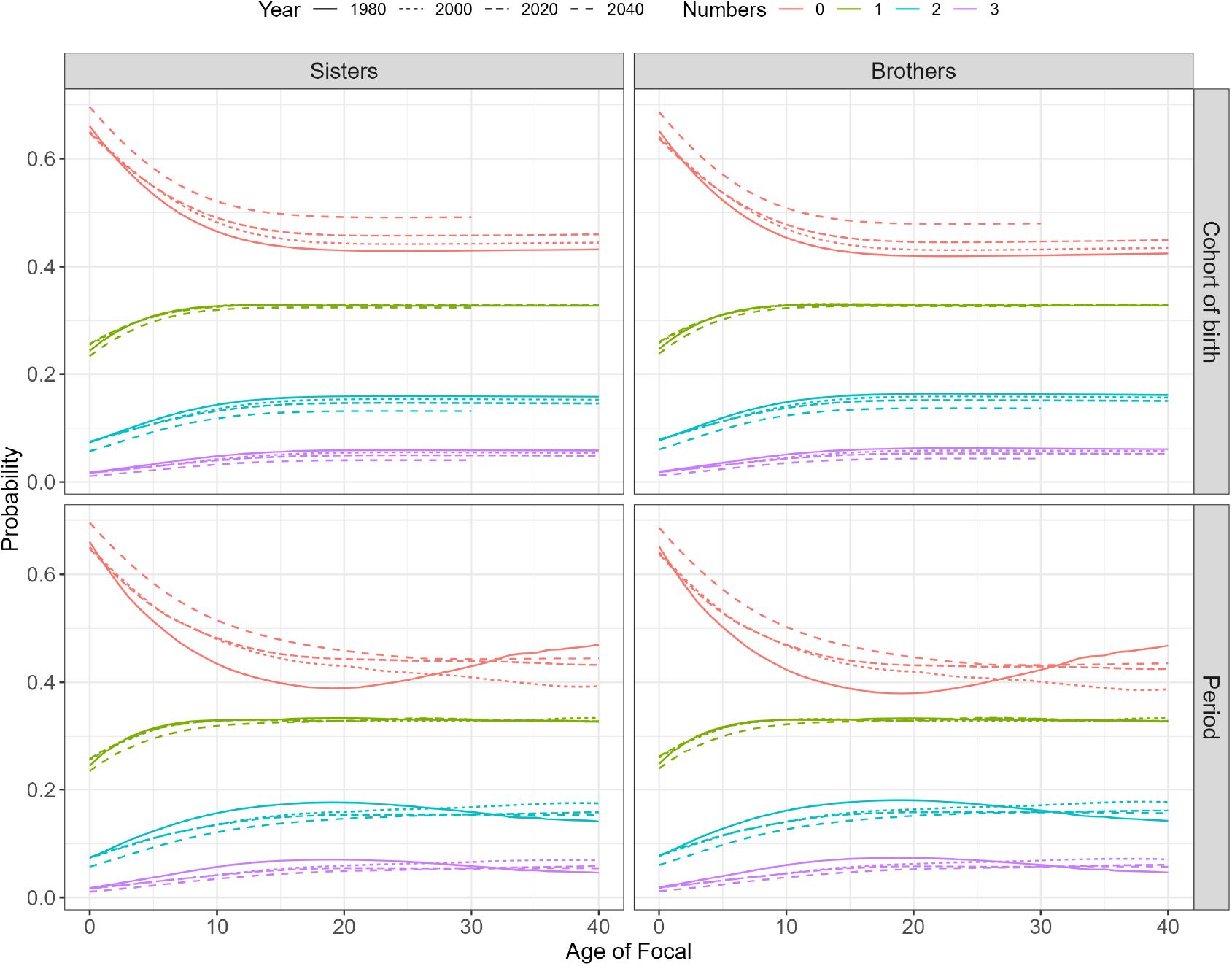
The probable numbers (colour) of sisters (left) and brothers (right) of Focal. Top: by age of Focals born to different cohorts. Bottom: by age of a typical Focal sampled from a given time period (here each age of Focal along the *x* represents a different cohort of Focals).

Fig 6 illustrates, for typical a Focal sampled from the years 2030, 2050, and 2070, the probable numbers of nieces, nephews, aunts, and uncles that she will have when she is aged 50. Compared to the other time periods, we see that in 2070, Focal (born in 2020) will have the greatest probability of having no nieces or nephews. This result is explained by declining fertility in our rate forecasts, which will result in Focal’s siblings having fewer probable children. Contrastingly, we predict Focal to have no uncles in 2050 with a greater probability than no uncles in 2070.

**Figure 6.**
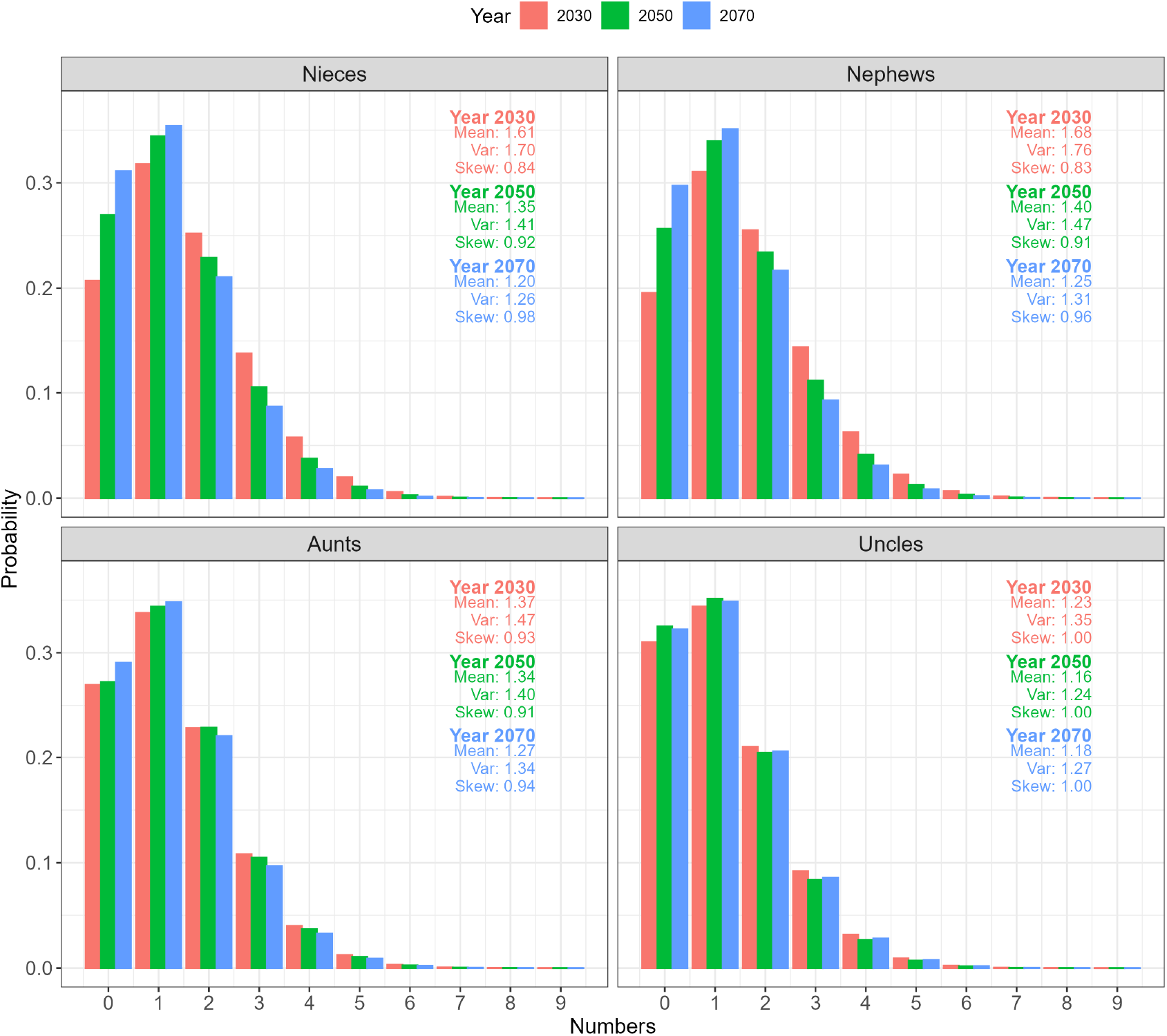
The probable numbers of nieces (top left), nephews (top right), aunts (bottom left) and uncles (bottom right) of Focal when Focal is aged 50. Colour represents time period from which we sample Focal, or equivalently, Focal’s cohort of birth, e.g., 2030 demonstrates a 50 year old Focal born in the 1980 cohort; 2070 illustrates a 50 year old Focal born in 2020.

### 3.4 Extension: the probable numbers of kin deaths

Fig 7 plots the probabilities that Focal loses *j* = 0, …, 3 siblings by the time she is 50. Results are presented from the perspective of typical Focals born to different cohorts (as indicated by the columns). The unconditional probability of losing one or more siblings includes the probability that the sibling was not born. Whereas, the conditional probability – through simple application of Bayes’ rule – conditions on one or more siblings being extant between Focal’s birth and her current age, but dying in the interim. We observe that the 1920 cohort (i.e., those aged in their mid 20s during WWII) experience considerably greater probabilities of experiencing a death of one or more sibling. This is likely explained by siblings being of similar age to Focal and therefore subject to the high war-time mortality rates.

**Figure 7.**
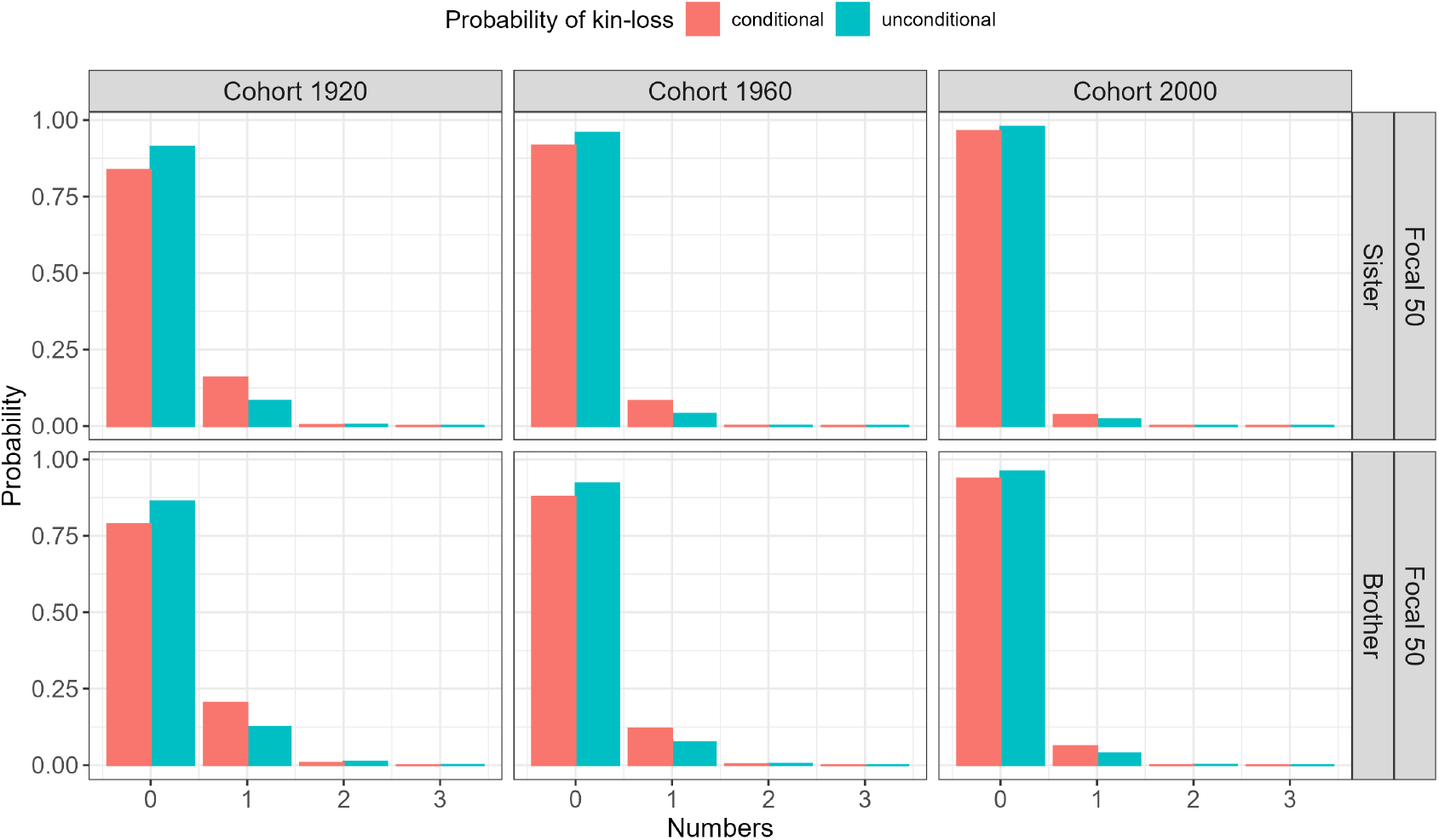
The conditional and unconditional probabilities of death of one or more sibling of Focal by her being aged 50. Note: *j* = 4+ probabilities not shown for visual aid.

In Fig 8 we plot the conditional probabilities that Focal experiences a bereavement of one or more sibling over her being aged 10 to 50. We compare these probabilities from the perspective of the cohort of birth of Focal.

**Figure 8.**
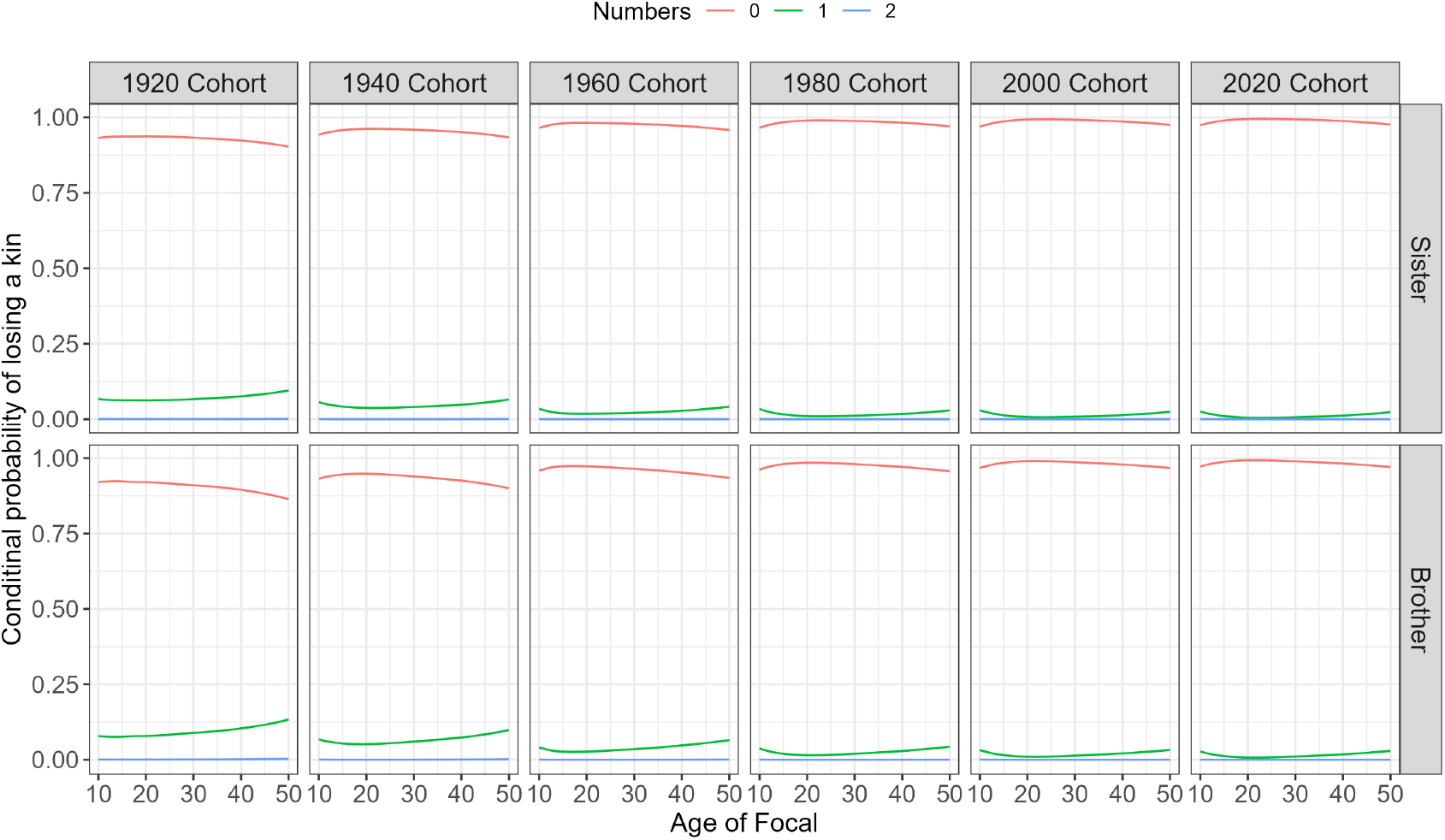
The conditional probabilities of death of one or more sibling of Focal by age of Focal 10 to 50. Note: *j* = 4+ probabilities not shown for visual aid.

## 4 Discussion

This research extends Butterick et al. (2025b) to the first kinship model able to predict the probable numbers of relatives, structured by age and sex within a time-varying demography. Formulae presented here are concise closed form expressions. They extend arbitrary genealogical distances to recover relatives considered in the leading frameworks of kinship (Alburez-Gutierrez et al., 2023; Caswell & Song, 2021; Caswell, 2022). They are able to reproduce the mean-field results presented therein (i.e., expected numbers of kin). As well as producing the probable numbers of living kin, the model flexibly extends to give the probable numbers of deaths an individual experiences. Such a detailed analysis of the kin-network will be useful in many fields.

Behind age-specific demographic rates are distributions; either assumed from probability theory, or from empirical data. By utilising these distributions corresponding to mortality and fertility schedules, rather than simply the rates, our framework produces a fully probabilistic representation of the kin-network. This contribution sets our framework apart from previous ones. One has flexibility to choose from any appropriate distribution to represent the random variable of births. For progress here we assume Poisson fertility. There are however no restrictions in presenting a more detailed analysis, e.g., using empirical distributions, derived from survey data or other sources.

Our approach allows one to consider kin-types of arbitrary relatedness, without requiring restrictive Markovian assumptions, or time-invariant population structures (Coste, 2025b). Formulae here, moreover, circumvent the need to incorporate complex time-and-generation-dependent multi-type branching processes to obtain kin-number distributions. This innovation considerably reduces the size of the state space required in the matrix projections. Although somewhat mathematically ugly, the extensive use of Fourier transforms and convolutions (implementable in all software) improves efficiency of the model.

The present model is restricted to age-structured populations. Recently, remarkable progress by Coste (2025b) has considered the expected numbers of kin of a typical individual, under any population structure. The inclusion of stage in population structure would be a valuable extension of the present work. One could then consider for instance, the probable numbers of kin structured by social environment, parity level, geographic location, or any generic life stage. In the current framework here, a multi-state extension would be complicated and computationally demanding; one would have calculate conditional Bernoulli/multinomial sequences, whereby, so many kin survive age-classes, and then transition stages. Working with the probability generating function as in the approach of Coste (2025a) and Tuljapurkar et al. (2020) could pave the way forwards. In any case, progress here would allow for a greater understanding of kinship structures. Stage-based models would allow one to condition fertility to depend on time since last birth (as well as age). This innovation would provide a more accurate description for human kinship, e.g., where time-proximity between births creates “sibling constellations” (Morosow & Kolk, 2017) and in other species which experience postpartum infertility (Ellis et al., 2024). Implementing stochastic age*×*stage structured kinship remains an open research problem.

Moving forwards, this research will be relevant in many areas. In age-structured animal populations, the model will be of use in estimating the probable numbers of specific kin, whereby kin are distanced by measures of relatedness. Such information will help in understanding the dynamics of social groups where post-reproductive relatives transfer vital information He et al. (2025). Regarding social policy in humans, we quantify variations in the numbers of relatives of individuals born to different cohorts. Knowing how these probable numbers responds to demographic trends, moreover, enables rate projections to be used to probabilistically predict future numbers of kin (Margolis & Wright, 2016). Applying the model to predict the probable numbers of bereavements that an individual experiences, will be particularly pertinent in understanding the social costs of family loss during conflicts. This analysis will directly supplement the research of Gómez-Ugarte et al. (2025), delimiting the mortality of humanitarian criss to the family level. Schlüter et al. (2023) used the framework of Caswell (2022) to estimate the number of children that experienced the death of a parent in the Gaza strip in 2023. The present model could extend this analysis to quantify the level of bereavement over all kin-types. Further application of the present framework could, for example, predict the number of relatives who require care (and/or have conditions), over age and time. Such progress will be of great interest in health and social care planning, and is currently ongoing.

## A Appendix Replenishment of kin

Here, we break down the equation in text (with *γ*_*o*_ ∈ {1, 2}; offspring sex, and *γ*_*p*_ ∈ {1, 2}; producer sex):

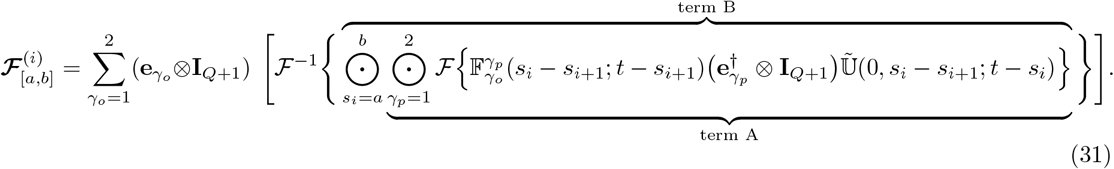

For arbitrary *i* > 0, consider the pmfs for the newborn *i*-th generation descendant of Focal’s *q*-th ancestor, 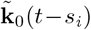. Suppose that these kin have offspring at exact age *s* = *s*_*i*_ −*s*_*i*+1_ (at time *t*−*s*_*i*+1_). At reproduction, the kin vector of pmfs is 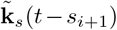. To be at reproductive age *s*, the kin survived that many years, starting from age 0 at time *t* − *s*_*i*_. That is, 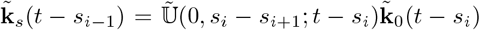. Recall that we encode female (male) kin by f = 1 (m = 2). As such, let *γ*_*p*_ = 1, *γ*_*p*_ = 2, *γ*_*o*_ = 1 and *γ*_*o*_ = 2. Consider for the case when *γ*_*o*_ = 1 (female offspring). Then, “term A” in the big square brackets reads:

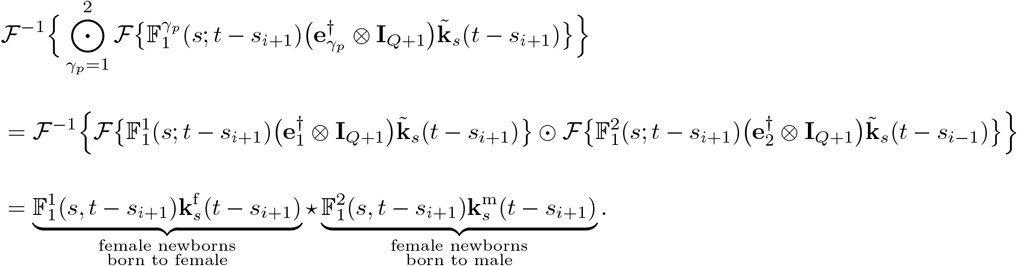

Next consider the case when *γ*_*o*_ = 2 (male newborns). Then, “term A” in the big square brackets becomes:

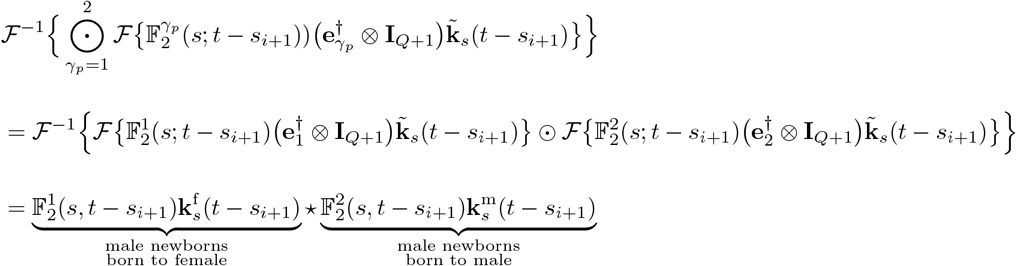

Continuing the above procedure over all possible fertile ages of reproducer kin, 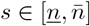 at times *t − s*_*i*+1_ + 1, *t − s*_*i*+1_ + 2, …, we arrive at “term B” in the square brackets:

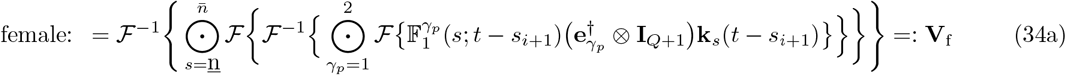

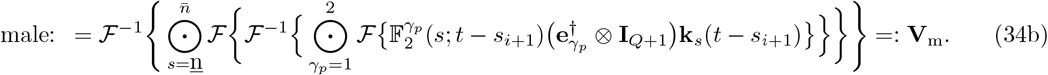

The action of the summand term 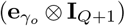 creates a block-structured vector: *γ*_*o*_ = 1 selects the term **V**_*f*_ padding the lower-block with zeros, *γ*_*o*_ = 2 selects **V**_*m*_ padding the upper-block with zeros, and adds them together:

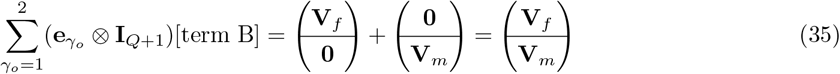

so that the pmf of offspring is in the correct block-structured form of females on top of males. Thus Eq (34a)-Eq (34a) are written in a block-structured form:

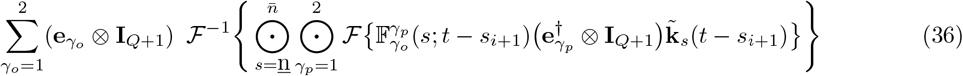

### B Appendix: Lines of ancestry

For instance, consider Focal’s female *i* = 1 and *i* = 2 generation ancestors. If *i* = 1 we have

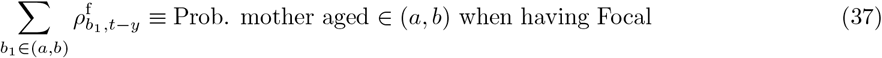

whereas when *i* = 2 we have two lineages:

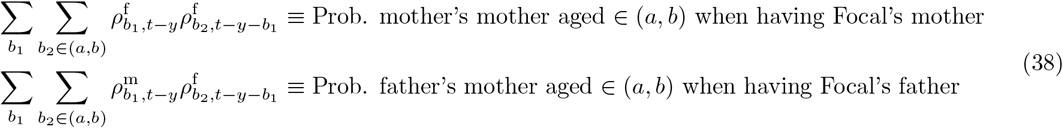

Note that for each specific sequence of ancestral events: [*b*_1_, *t − y*], [*b*_2_, *t − y − b*_1_], …, [*b*_*q−*1_, *t − β*_*q−*2_], we have that

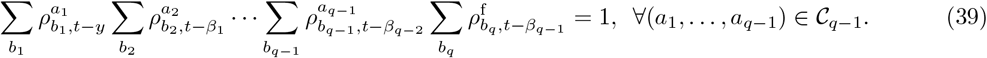

### C The probable number of kin deaths

Consider the probable number of kin of sex *γ* that die between being age *s*′ at time *t*, and prospective age *s*

(if survived). From a little manipulation of Eq (8) in text, we can construct a matrix 𝔻^*γ*^ (*s*′, *s*; *t*) with (*i, j*) entries given through:

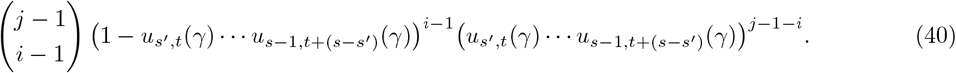

Over both sexes we build 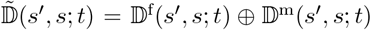. Now consider the probable numbers of kin which die between Focal being born and her being age *y*.

There are two cases to consider depending on whether *s*_*g*_ > *y* or *s*_*g*_ < *y*: If the former, then

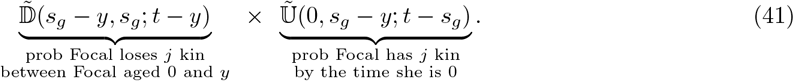

while if *s*_*g*_ < *y* then

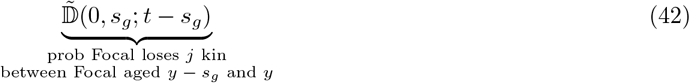

Letting *z* = max*{*0, *s*_*g*_ *− y}*, and recalling by definition that 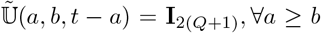 and 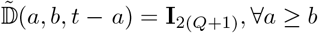, we simply write 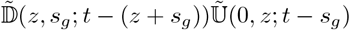

#### C.1 The basic logic

Eq (41)-Eq (42) provide the unconditional probabilities that Focal experiences so many kin deaths, including the case in which no kin are born (adding to the zero entry). Consider Eq (42), restrict to female kin, and omit sex and time notations. Recalling that **k**_*s*_ = (*k*_*s*_(0), …, *k*_*s*_(*Q*))^*†*^ is the pmf of kin at age *s*, let 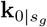 represent newborn kin who will be age *s*_*g*_ when Focal is *y*. Then

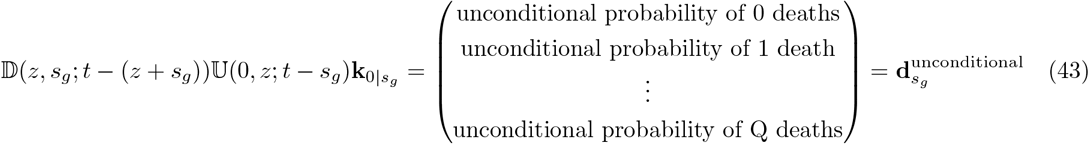

has entries: the total probability that Focal experiences *j* deaths of kin who would be age *s*_*g*_ when she is aged *y*, between her being aged zero and age *y*. Over the age-range *s*_*g*_ ∈ Σ the total probability can be written as the (*j −* 1)-th entry of the convolution:

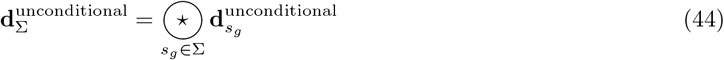

We might however want the conditional probabilities that *j* kin die, given there is at least one kin extant (i.e., kin who are born). Again, restrict to female kin, and omit sex and time notations. Let this vector be **d**^conditional^. We apply Bayes’ formula to derive the conditional probabilities:

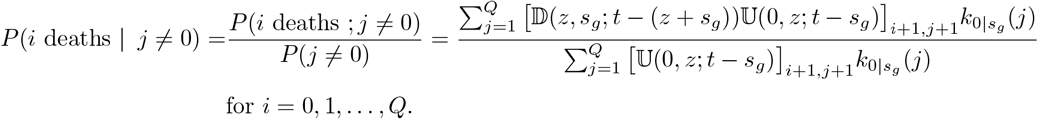

Again let the pmf of newborn kin who would be age *s*_*g*_ when Focal is *y* be 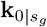. In matrix form:

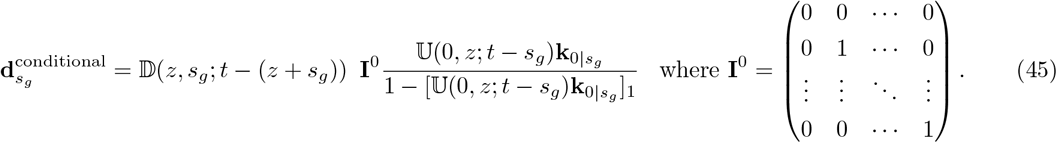

Here, if *z* = 0 (*s*_*g*_ < *y*) we have the conditional pmf of newborns (conditioned on *j* > 0), 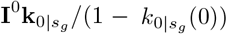. The probable numbers of deaths of these kin up to Focal *y* is given by the pre-multiplication by 𝔻(0, *s*_*g*_; *t − s*_*g*_). If *z* = *s*_*g*_ *− y* (i.e., *s*_*g*_ > *y*) we have the conditional pmf of kin at Focal’s birth, 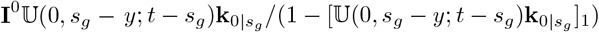. Pre-multiplication by 𝔻(*s*_*g*_ *− y, s*_*g*_; *t − y*) gives the probable number of deaths up to Focal being *y*.

Over the age-range *s*_*g*_ ∈ Σ the total conditional probability is written as as a weighted sum:

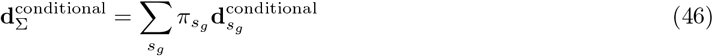

where 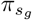 is the conditional probability that given *j* > 0 kin are alive at birth, the kin is now aged *s*_*g*_ when Focal is age *y*:

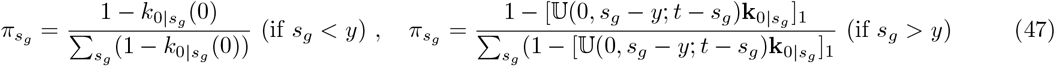

We apply the above to pmfs of newborn kin who would be age *s*_*g*_ when Focal is *y*, obtained from Eq (21), Eq (24), Eq (27), and Eq (30). This allows us to derive the unconditional and conditional probabilities of deaths. Appendix C.2 provides a general example of kin structured by lineage type, and Appendix C.4 provides an explicit example of siblings, used in the Application section in text.

#### C.2 An example: kin which descend through younger lineages

As an explicit example of the above, consider kin which descend through younger siblings of Focal’s (*q −* 1)-th ancestor, Eq (21). Recall we use *z* = max*{*0, *s*_*g*_ *− y}*.

##### C.2.1 Unconditional probabilities of kin-loss

The unconditional numbers of deaths of kin are found by simply substituting 𝔻 for 𝕌:

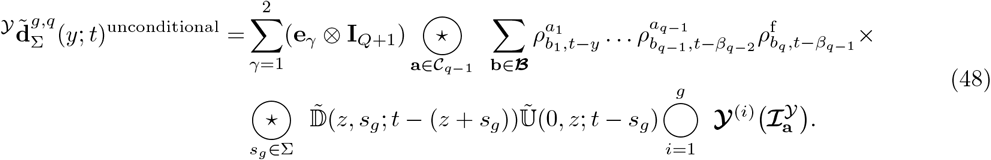

##### C.2.2 Conditional probabilities of kin-loss

The conditional numbers of deaths of kin are more complex to derive. We find the probable numbers of kin-loss to be given by:

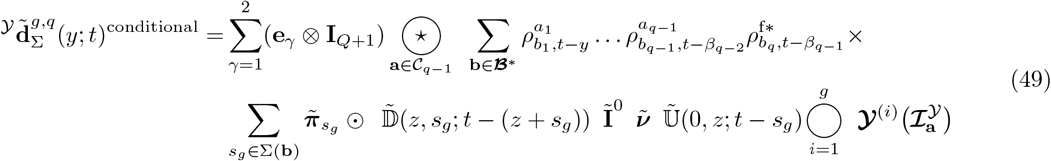

Here the set **ℬ**^∗^ restricts the ancestral ages to rule out the possibility of Focal’s *q*-th ancestor “not” reproducing a younger sibling of her (*q −* 1)-th ancestor (we must condition on birth of the chain of descent):

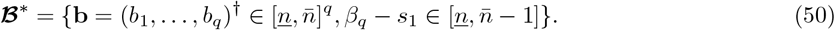

The set **ℬ**^∗^ thus defines conditional probabilities 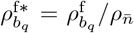 used in the summation over ancestral ages.

Additionally, the possible age-ranges of kin of age *s*_*g*_ are restricted to:

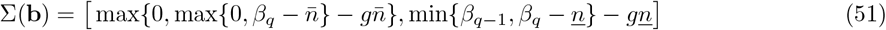

See Appendix C.5 for derivation of the constraints. For notational ease, we set

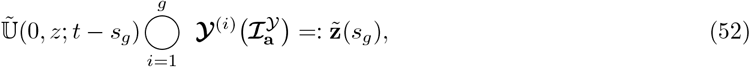

where 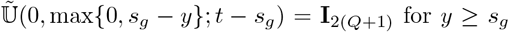. Thus, we see that 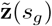 either represents (i): the kin-number pmf of kin at birth of Focal (if *y* < *s*_*g*_) or (ii): the pmf for newborn kin when Focal is aged *y − s*_*g*_ (if *y* > *s*_*g*_). For each kin of age *s*_*g*_ ∈ Σ(**b**), Eq (49) conditions on the (*g −* 1)-th descendant of Focal’s *q*-th ancestor having *j* > 0 offspring. That is we are applying a weighted sum of conditional pmfs, where each conditional pmf yields the probabilities that Focal experiences *j* deaths of kin who are born and would be aged *s*_*g*_ when Focal is *y*. As per Eq (45), to calculate each conditional pmf, we rule out the possibility of 0 extant (born) kin i.e.,

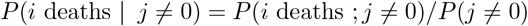

using:

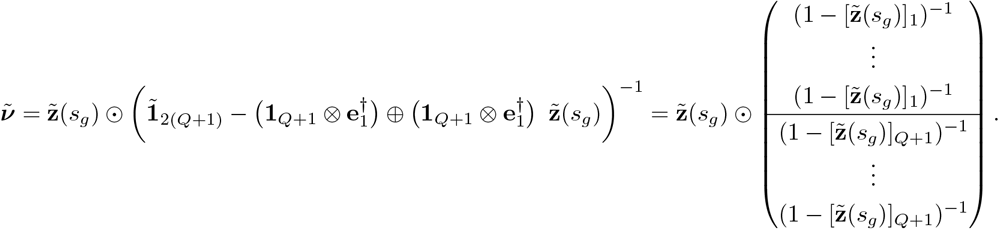

The variable and 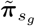 weights the conditional probabilities, on the event “at least one kin is extant by age *s*_*g*_ *− y*”, over all possible ages the kin could be; *s*_*g*_ ∈ Σ(**b**):

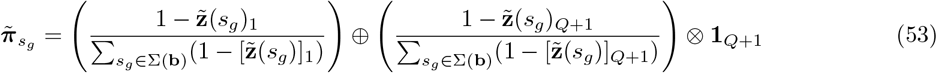

**To summarise: for each conditional b** ∈ ℬ^∗^, **we condition on at least one kin being born and calculate the conditional probability mass function that the kin will be of age** *s*_*g*_ **when Focal is** *y*. **We then probabilistically sum over all of possible ages of kin** *s*_*g*_ ∈ Σ(**b**) **with appropriate weights (still conditioned on ancestor ages). We then sum over the feasible set of parental ages** ℬ^∗^ **which strictly allows for the chain of events to procure a younger lineage**.

#### C.3 Combining deaths from younger and older lineages

Here, we seek the overall conditional pmf, *d*_total_(*j*), defined as the probability that Focal has lost *j* kin of particular sex, conditional on Focal having at least one kin (either through the younger or older lineage), (*l*_total_ > 0):

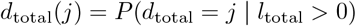

For notational ease we remove sex and time notation here. Let

**p**_*Y*_ = *p*_*Y*_ (*j*) and **p**_*O*_ = *p*_*O*_(*j*) be pmfs for the number *j* = 0, …, *Q* of younger and older kin of Focal extant between Focal’s birth and age *y*.

**d**_*Y*_ = *d*_*Y*_ (*j*) be the pmf for the number (*j* = 0, …, *Q*) of lost younger kin conditional on kin being extant between Focal’s birth and age *y* (i.e., Eq (49)).

**d**_*O*_ = *d*_*O*_(*j*) be the pmf for the number (*j* = 0, …, *Q*) of lost older kin conditional on the kin being extant between Focal’s birth and age *y* (i.e., Eq (49)).

The probability space is partitioned into exclusive events: *ω*_1_, *ω*_2_, *ω*_3_. Let *l*_*Y*_ represent the number of younger lineages of Focal extant between Focal’s birth and age *y*, and *l*_*O*_ the same for older.

1. *ω*_1_: *l*_*Y*_ > 0 **and** *l*_*O*_ = 0. As such, the total deaths *d*_total_ = *d*_*Y*_ and *ω*_1_(*j*) = *d*_*Y*_ (*j*)
2. *ω*_2_: *l*_*Y*_ = 0 **and** *l*_*O*_ > 0. As such, the total deaths *d*_total_ = *d*_*O*_ and *ω*_2_(*j*) = *d*_*O*_(*j*)
3. *ω*_3_: *l*_*Y*_ > 0 **and** *l*_*O*_ > 0. Here, the total deaths *d*_total_ = *d*_*Y*_ + *d*_*O*_ (conditional independence ⇒ the convolution of *d*_*Y*_ and *d*_*O*_, and as such, *ω*_3_(*j*) = (*d*_*Y*_ *\* d*_*O*_)[*j*]

The above events are weighted by *w*_*c*_; the probability of event *ω*_*c*_ occurring, conditional on the overall event *l*_total_ > 0. That is, *w*_*c*_ = *P* (*ω*_*c*_ | *l*_total_ > 0). The probability of the overall condition is the complement of Focal experiencing zero of the kin:

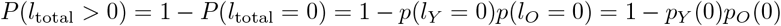

Then using independence:

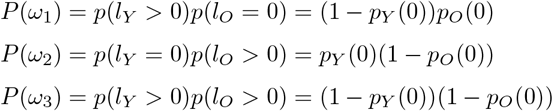

where

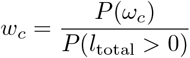

the overall conditional pmf *d*_total_(*j*) is the weighted average (law of total probability) of the three pmfs:

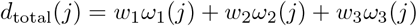

or in vector form:

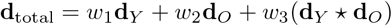

### C.4 Explicit example of siblings

The kin-number pmf for younger siblings of age *s*_1_ ∈ Σ = [0, *y* − 1] is

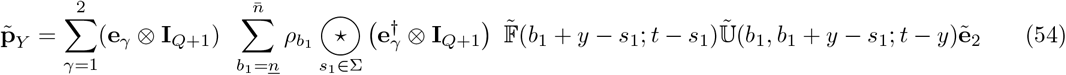

The conditional probabilities of Focal experiencing at least one younger sibling restricts the set 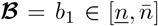 to 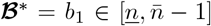 with conditional probabilities defined by 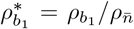 for 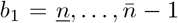. Each conditioning *b*_1_ restricts the set 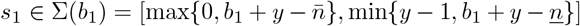, e.g., if Focal is *y* = 10 and 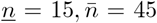, if *b*_1_ = 15, 16, …, 35, 36, …, 44 then younger sisters of Focal can be of ages [0, 9], [0, 9], …, [0, 9], [1, 9], …, [9, 9]. The pmf for the probable numbers of deaths, conditional on a younger sister being born is

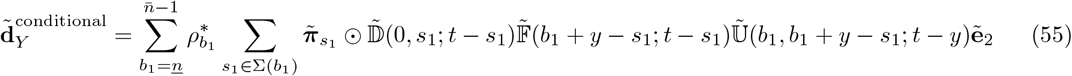

where for each conditioning *b*_1_, we have that:

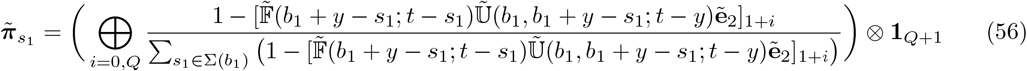

For older siblings we have that

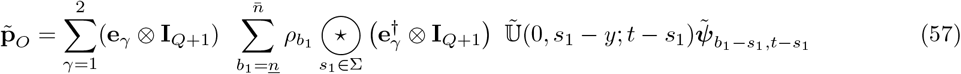

In this case, the conditional probabilities of Focal experiencing at least one older sibling restricts the set 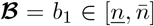 to 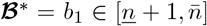 with conditional probabilities defined by 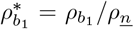 for 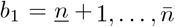. Each conditioning *b*_1_ restricts the set 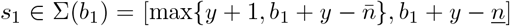. The pmf for the probable numbers of deaths, conditional on a older sister being born is:

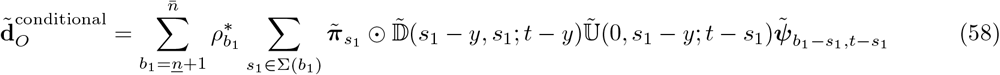

where for each conditioning *b*_1_, we have that:

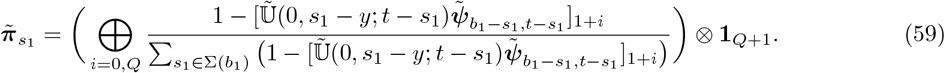

### C.5 Constraints

The following three inequalities need to be satisfied for kin descending through younger and older lineages:

1. **Ancestral ages**: 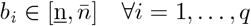
2. **Ages of descendant reproduction**: 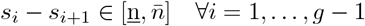
3. **Age of the** *q***-th ancestor** when producing descendant (now aged *s*_1_):

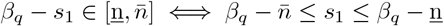

#### C.5.1 Constraints (kin through younger lineages)

In addition to the above conditions, here we have that:

1Y. **Sibling of** (*q* − 1)**-th ancestor younger**:

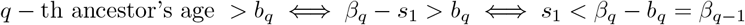

Current age of kin, *s*_*g*_, is constrained by the bounds on the age *s*_1_. Let 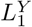 and 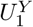 be the lower and upper bounds of *s*_1_ derived from 1,2,3, and 1Y:

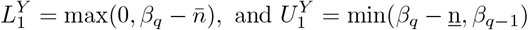

meaning that *s*_*g*_ is constrained by 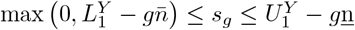.

#### C.5.2 Constraints (kin through older lineages)

An additional inequality here is:

1O. **Sibling of** (*q* − 1)**-th ancestor older**:

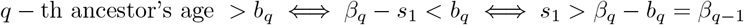

Age of kin *s*_*g*_ is again constrained by *s*_1_. Let 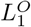 and 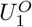 be the lower and upper bounds of *s*_1_, where using 1,2,3, and 1O:

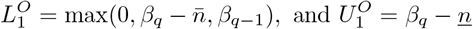

and as such, we have 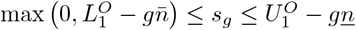.

## Footnotes

1 that an individual will perform an altruistic act to another if the coefficient of relatedness between the two exceeds the cost-to-benefit ratio of performing the act

